# Genetic ablation of dentate hilar somatostatin-positive GABAergic interneurons is sufficient to induce cognitive impairment

**DOI:** 10.1101/2022.11.01.514756

**Authors:** Rajasekar Nagarajan, Jinrui Lyu, Maltesh Kambali, Muxiao Wang, Connor D. Courtney, Catherine A. Christian-Hinman, Uwe Rudolph

## Abstract

Aging is often associated with a decline in cognitive function. A reduction in the number of somatostatin-positive (SOM+) interneurons in the dentate gyrus (DG) has been described in cognitively impaired but not in unimpaired aged rodents. However, it remains unclear whether the reduction in SOM+ interneurons in the DG hilus is causal for age-related cognitive dysfunction. We hypothesized that hilar SOM+ interneurons play an essential role in maintaining cognitive function and that a reduction in the number of hilar SOM+ interneurons might be sufficient to induce cognitive dysfunction. Hilar SOM+ interneurons were ablated by expressing a diphtheria toxin transgene specifically in these interneurons, which resulted in a reduction in the number of SOM+/GAD-67+ neurons and dendritic spine density in the DG. C-fos and Iba-1 immunostainings were increased in DG and CA3 but not in CA1. Behavioral testing revealed a reduced recognition index in the novel object recognition test, a reduction in the percentage of correct alternations in the Y maze tests, and increased latencies and path lengths in the learning and the reversal learning phase of the Morris water maze. Our results show that partial genetic ablation of SOM+ hilar interneurons is sufficient to increase activity in DG and CA3, as has been described to occur with aging and to induce an impairment of learning and memory functions. Thus, partial ablation of hilar SOM+ interneurons may be a significant contributing factor to age-related cognitive dysfunction. These mice may also be useful as a cellularly defined model of hippocampal aging.

## Introduction

Normal cognitive aging is characterized by accelerated declines in memory from early adulthood[1], and the factors behind this process are not entirely clear. Increasing age is a risk factor for postoperative cognitive deficits, which were previously called postoperative cognitive dysfunction (POCD)[2, 3] and are now referred to as postoperative neurocognitive disorder (poNCD)[4]. Postoperative cognitive deficits can persist for many months after surgery, e.g., in patients > 60 years of age, approximately 10% of patients displayed cognitive dysfunction three months after surgery, compared to 3% in a control group[5]. The molecular and cellular mechanisms underlying cognitive impairments in older individuals remain poorly defined. Mounting evidence points to changes in GABAergic function as a major causal factor. Indeed, GABA levels in frontal and in parietal areas decrease with age[6], and GABA levels are substantially reduced in hippocampus, striatum, and cortex in aged mice (18 months)[7]. Such declines in GABA levels may suggest that functional deficits in the GABA system contribute to age-related cognitive dysfunction. Moreover, in human postmortem studies it has been found that the expression of the GABA_A_ receptor α5 subunit, which has previously been shown to play an important role in modulating learning and memory functions in the hippocampus [8, 9], is decreased in aged human individuals[10]. Decreased GABA neurotransmitter levels were also observed in the CSF of healthy humans with aging[11].

One of the most studied brain structures affected by aging is the hippocampus, which plays an important role in learning and in the formation and consolidation of memory[12]. During aging, the hippocampus shows a decrease in volume, which correlates with a decline in learning and memory[13]. Importantly, GABAergic interneurons have been linked to memory processes. Optogenetic inhibition of hilar GABAergic interneuron activity impairs spatial memory[14]. An immunohistochemical study analyzing interneurons in different hippocampal subregions revealed that in aged memory-impaired rats the number of SOM+ interneurons in the DG hilus are significantly smaller than in age-unimpaired rats[15]. Recent studies in humans and rodents have shown that the different hippocampal subregions are differentially affected by aging. In human subjects, older individuals have increased activity in CA3/DG during while performing pattern separation tasks[16], and in rodents, aging-related dysfunction is associated with hyperexcitability of CA3 pyramidal cells but hypo-excitability of CA1 pyramidal cells [17]. Causal relationships between dysfunction in DG, CA3, and CA1 are unclear, but it has been speculated that hypo-excitability of CA1 pyramidal cells may occur in compensation for DG/CA3 hyperactivity, as the DG/CA3 hyperactivity precedes changes in CA1[16]. Furthermore, Koh et al. (2013) reported that selective positive allosteric modulation of α5GABA_A_ receptors improves cognitive function in aged rats with memory impairment [18].

It has been shown that α5-GABA_A_ receptor-mediated tonic inhibition in the DG plays an important role in ensuring normal cognitive functioning under high-interference memory conditions[9]. Recent evidence supports the cognitive salience of inhibitory interneurons and their role in maintaining balanced excitation-inhibition in the aging human brain [16, 19], and studies have shown interneurons to be specifically vulnerable in brain aging [15, 20]. Fast spiking interneuron deficits and secondary alterations in high frequency gamma cortical network oscillations were noted in aged mice, the latter being fundamental to higher order information processing[19, 21]. Reductions of SOM-immunoreactive interneurons were observed in the temporal cortex in AD patients[22]. In addition, age-related cognitive deficits have been found to be correlated with a decrease in SOM-positive interneurons in the DG hilus in rodents [15, 23]. However, it remains unclear whether the reduction in SOM+ interneurons in the DG hilus is directly related to or even causal for age-related cognitive dysfunction.

To understand why elderly individuals are more prone to poNCD than younger ones, it will be helpful to better understand the mechanisms that contribute to decreased cognitive function in the elderly. Therefore, we studied the role of a reduced number of somatostatin-positive interneurons in the DG hilus for age-related cognitive deficits. We hypothesized that hilar (SOM+) interneurons play an essential role in maintaining cognitive function and that a reduction in the number of hilar (SOM+) interneurons might be sufficient to induce cognitive dysfunction. To test our hypothesis that loss of numbers of these interneurons underlies age related cognitive dysfunction at least in part, we ablated SOM+ interneurons in the DG hilus.

## Materials and Methods

### Animals

Adult Sst-IRES-Cre transgenic mice (Stock no. 013044, The Jackson Laboratory) were crossed with C57BL/6J mice (Stock no. 000664, The Jackson Laboratory). Hemizygous adult mice of both sexes (3-4 months) were used for all studies. To determine whether the cellularly defined model (ablation of Hilar SOM+ interneurons) has any behavioral features of age-related cognitive dysfunction, we performed behavioral experiments with two control groups [young adult (3-4 months) and aged (18-19 months) C57BL/6J wild type mice]. 8 to 12 weeks old C57BL/6J wild type mice were obtained from The Jackson Laboratory (Bar Harbor, Maine). The mice were housed in groups of 4–5 per cage. Mice that showed aggression towards cage mates were singly housed. The animals were monitored for health on a weekly basis but otherwise left undisturbed until behavioral characterization at 3-4 months old (young) or 18–19 months old (old or aged). All mice were housed in a climate-controlled room (temperature: 22 ± 2°C; humidity: 55 ± 5%) and maintained on a 12:12 light-dark cycle (0700-1900) with food and water ad libitum. The maintenance and handling of mice were consistent with the Guide for the Care and Use of Laboratory Animals of the National Research Council. All procedures were approved by the Institutional Animal Care and Use Committee of the University of Illinois Urbana-Champaign.

### Stereotactic surgery

To generate mice with a partial ablation of the SOM+ interneurons in the dentate hilus, AAV5-EF1α-mCherry-flex-dtA (construct-387-aav2-5 from the viral vector core of the Canadian Neurophotonics Platform) was injected bilaterally into the DG hilus of Sst-IRES-Cre mice (3-4 months of age). Mice received ketoprofen 5mg/kg, s.c., and atropine 0.04 mg/kg, s.c. before surgery. Mice were anesthetized with isoflurane (3%) and maintained under anesthesia (2 %) throughout the surgery. The AAV vector was injected bilaterally into the DG hilus (stereotactic coordinates AP: 2.1mm, ML: ±1.5mm, DV: −2.1mm relative to bregma) at a rate of 120 nl/min (1,000 nl per side), and the needle was left in place for an additional 8 mins to permit diffusion. The control groups were injected with an AAV5-EF1α-mCherry virus (VN2098 from Vector Biolabs). All surgical procedures were performed under aseptic conditions. Mice were kept warm on a heating pad as they recovered. Mouse chow moistened with water was placed in the cage once they recovered to encourage eating and provide hydration following surgery. To allow time for recovery and viral expression, animals were housed for three weeks after injection and before any behavioral experiments were initiated. Injection locations were verified histologically at the end of the study.

### Behavioral Tests

All behavioral tests were performed in the morning, with at least 1 h of habituation in the behavioral room prior to experimentation on each day.

### Novel Object Recognition Test

Mice were habituated in the experimental chamber for two days before the testing phase, and each habituation period lasted for 15 min for each animal per session. During the training phase on day three, the mouse was allowed to explore two copies of the same object for 10 min. The mouse was then returned to its home cage for a retention period of 1 h. The mouse was reintroduced to the training context and presented with one familiar sample object and one novel object for 10 min. Movement and interaction with the objects were recorded with the EthoVision XT video-tracking system, with a videocamera mounted above the chamber, and exploratory behavior was measured. Interaction time was recorded using the multiple body point module. Memory was assessed using the recognition index (i.e., the ratio of time spent exploring the novel object to the time spent exploring both objects). After each mouse had performed the test, the chamber and objects were swabbed thoroughly with alcohol to avoid any interference with tests involving subsequent mice.

### Morris Water Maze and Reversal Learning

The spatial learning and memory function of rats were assessed using the Morris water maze. The test lasted for two weeks: Learning phase; Learning phase probe trials; Reversal learning phase & Reversal learning probe trials. Mice are tested in a pool (120 cm in diameter) filled with water (23-25°C) made opaque with the addition of a white nontoxic dye (Premium Grade Tempera, Blick, Galesburg, IL) containing a platform (10 cm in diameter) that is submerged by 1 cm under water surface. Geometric shapes are affixed to the walls to serve as extra-maze cues. Mice are given 3 trials every day released from a different quadrant each time in random order, with the platform location constant. A trial ends either 2 sec after the mouse climbs on the platform, or 60 sec after the start of the trial, with the experimenter guiding the mouse to the platform. If a mouse could not find the platform within 60 sec it was gently guided to the platform and the escape latency was recorded as 60 sec. After the mouse reached the platform, it was allowed to stay there for 10 seconds before the next trial was performed. The reversal learning phase was established by moving the platform from the original location to the nearest quadrant to increase the effects of interference. During probe trial and reversal learning probe trial, the platform was removed, and the mice were left in the pool for 60 seconds. The track of the mice was recorded using a camera positioned above the center of the pool and connected to the EthoVision XT software video-tracking system.

### Y-maze test

Spontaneous alternation in the Y maze was tested as a measure of spatial working memory. The apparatus was a Y-shaped maze with three gray, opaque plastic arms at a 120° angle from each other. The arm dimensions were 15 cm (height) x 30 cm (length) x 8 cm (width). Each animal was placed in the center of a symmetrical Y-maze and was allowed to explore freely during an 8-min session. The sequence and total number of arms entered were recorded with a video camera that was mounted above the apparatus. Arm entry was defined as entry of the whole body into an arm. Alternation was defined as successive entries (ABC, ACB, BCA, BAC, CBA, CAB) into the three arms in overlapping triple sets. The % of alternation was calculated as the number of triads containing entries into all three arms divided by the maximum possible number of alternations (the total number of arms entered − 1) × 100. To diminish odor cues, the Y maze was cleaned with 70% ethanol between trials and allowed to dry.

### Biochemical and molecular parameters Immunofluorescence staining

Mice were deeply anesthetized with ketamine (139 mg/kg i.p.) / xylazine (21 mg/kg i.p.) and then transcardially perfused with ice-cold phosphate-buffered saline (PBS) followed by 4% ice-cold paraformaldehyde. After 24 hours of post-fixation at 4°C, brains were moved into 30% sucrose for 72 hours and sectioned into 30 µm coronal sections using a cryostat. Free-floating sections were collected mounted on slides for staining.

Immunofluorescence staining was used to validate hilar SOM+ interneuron manipulation. Free-floating coronal sections (30 μm) were nuclei stained with DAPI for 10 min in the dark, and the sections were transferred to glass slides and mounted with Anti-Fade Fluorescence Mounting Medium (Vector Laboratories). Images were acquired with a Leica microscope (Leica, Germany).

### Immunohistochemistry

We performed analysis of sections for target cell labeling and quantification. For analysis of c-fos staining, all mice (n=4 per group) were exposed to a novel environment for 60 min and then placed into a clean cage individually for 1 hour before perfusion. Free-floating coronal sections (30 μm) were incubated in blocking solution (2% normal goat serum and 0.4% Triton X-100 in PBS) for 2 h at room temperature and then with primary antibodies at 4°C overnight. The primary antibodies used were as follows: rabbit anti-c-fos (Cell Signaling Tech, cat # 2250, 1:500), anti-somatostatin (SOM, Invitrogen, cat # PA5-82678, 1:1000), anti-glutamic acid decarboxylase 67 (GAD-67, Invitrogen, cat # PA5-21397, 1:500) and anti-Iba-1 on microglia (Iba-1, Invitrogen, cat # MA5-36257, 1:500). After washing with blocking solution, we added 3% H2O2 for 10 min at room temperature. After washing with blocking solution, the sections were incubated in a biotinylated goat anti-rabbit secondary antibody in blocking solution (Invitrogen, cat # 31820, 1:500) and transferred in a detection reagent (Vectastain Elite ABC kit, Vector Laboratories, cat # PK-7200). Immunoreactivity was visualized using the VECTASTAIN ABC kit (Vector) for 30 min and Vector NovaRED (Vector) as the chromogen for 3 min at room temperature. Finally, brain sections were washed with distilled water, transferred to glass slides, dehydrated in 95% and 100% ethanol for 1 min each and mounted with Eukitt mounting medium (Sigma-Aldrich).

### Western blots

Mice were deeply anesthetized with ketamine (139 mg/kg i.p.) / xylazine (21 mg/kg i.p.) and then transcardially perfused with ice-cold normal saline. Brains were harvested from mice. Hippocampal tissue was homogenized in RIPA buffer containing protease inhibitors and phosphatase inhibitors. The homogenate was centrifuged at 13,000 rpm for 30 min at 4°C, and supernatant was collected for measurement of protein concentrations with a BCA assay kit. Proteins were separated by sodium dodecyl sulfate polyacrylamide gel electrophoresis (SDS-PAGE), transferred onto polyvinylidene fluoride (PVDF) membranes, and incubated at 4°C overnight with primary antibodies for Iba-1 [1:500, Invitrogen, catalog # MA5-36257] and β-actin [1:2,000, Cell Signaling Technology, catalog # 4970]. The membranes were washed with TBST three times and then incubated at room temperature for 60 min with secondary antibody goat anti-rabbit IgG (1:3000, Cell Signaling Technology, catalog # 7074). Then the membranes were washed with TBST four times. The SuperSignal West Pico Chemiluminescent Substrate kit from Thermo Fischer Scientific (Rockford, IL) was used to detect antigens utilizing the manufacturer’s instructions [24].

### Golgi-Cox Staining

Mice were deeply anesthetized with ketamine (139 mg/kg i.p.) / xylazine (21 mg/kg i.p.) and then transcardially perfused with ice-cold normal saline. Brains were harvested from mice and stained using the superGolgi Kit (Cat#003010, Bioenno Tech) according to the manufacturer’s instructions. We examined the spines of the dorsal dentate gyrus. Primary spines were not analyzed; secondary and tertiary dendrites were selected for analysis. Z-stacks of Golgi-stained dendrites were taken at 100 x magnification on a Kyence All-in-One Fluorescence Microscope (BZ-X 810). One segment per individual dendritic branch and two branches per neuron were chosen for the analysis. Spine densities per 20 μm calculated using Reconstruct 11 software (Version 1.1.0.1)[25].

### Statistical Analysis

Statistical analyses were performed (GraphPad Prism) using Two-way ANOVA followed by Sidak’s multiple comparisons test for multiple comparisons or two-tailed t-test. Results are represented as mean ± S.E.M. and p values are specified in each figure legend. p values less than 0.05 were considered significant.

## Results

In order to test whether a partial ablation of SOM+ interneurons induce an aging-related phenotype, we stereotactically injected AAV5-EF1-α-mCherry-flex-dtA into the DG hilus of Sst-IRES-Cre mice and started 21 days later with behavioral experiments, followed by molecular analysis (Fig. 1A). To validate the specificity of the hilar SOM+ interneuron manipulation, we performed immunofluorescence staining of coronal sections from experimental mice with DAPI counterstain and mCherry stain (Fig.1C). With the viral vector used, the mCherry expression should occur in every transduced cell in which the EF1-α promoter is active and independently of Cre-loxP-mediated recombination, whereas the expression of the diphtheria toxin subunit A is dependent on Cre-loxP-mediated recombination.

**Fig. 1.**
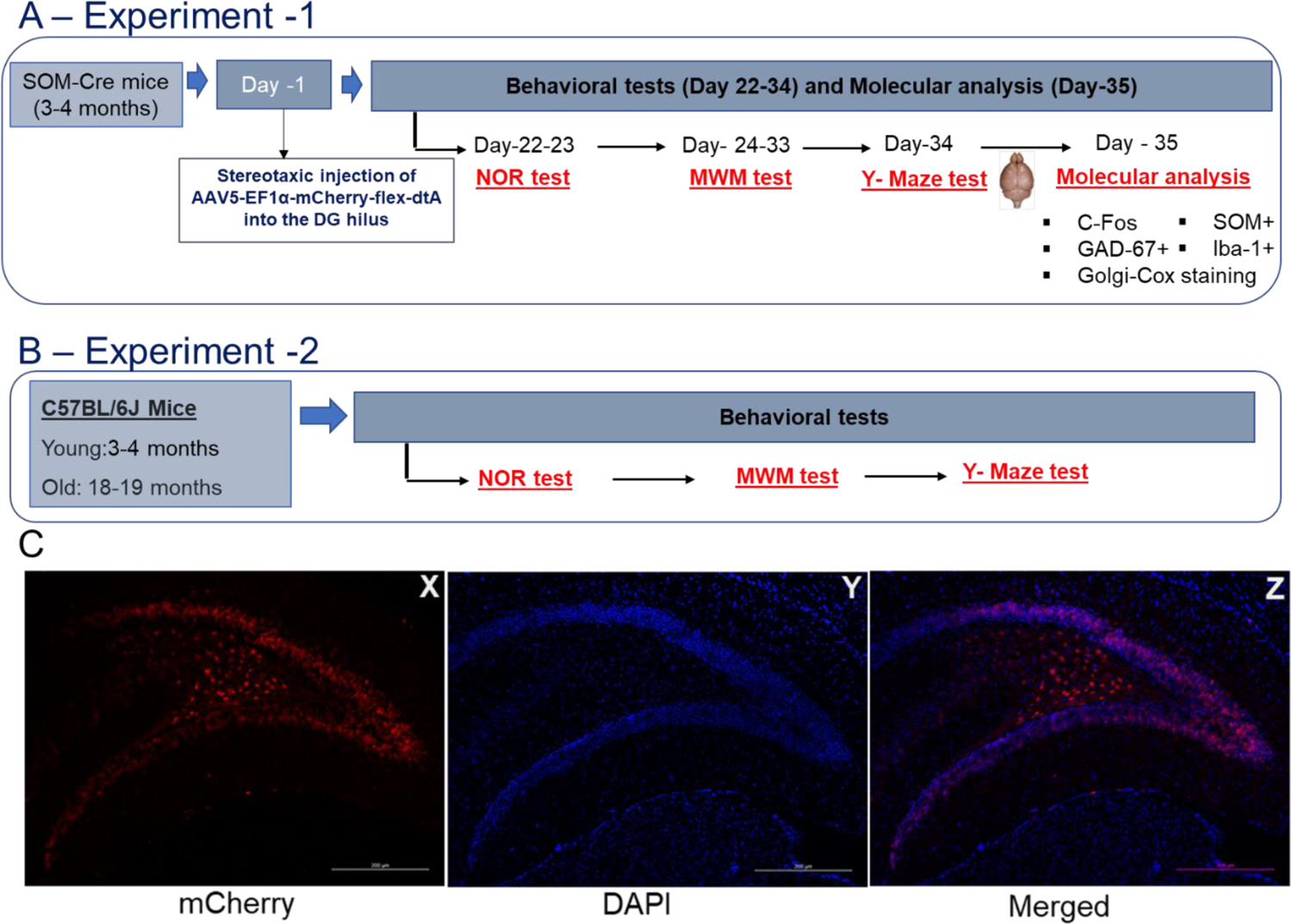
A&B) Schematic overview of the experimental protocol. A) Experiment 1: Testing whether genetic ablation of somatostatin-positive hilar interneurons results in cognitive deficits. B) Experiment 2: To determine whether the cellularly defined model (ablation of Hilar SOM+ interneurons) has any behavioral features of age-related cognitive dysfunction, we perform behavioral experiments with two control groups [young adult (3-4 months) and aged (18-19 months) C57BL/6J wild type mice]. C) Validation of viral injection into the DG hilus. The viral vector rAAV5/EF1-Lox-Cherry-lox(dtA)-lox-2.ape was injected into the DG hilus of SOM-Cre mice. The mice were perfused. Slices were stained with DAPI. The red mCherry fluorescence is shown in X, DAPI staining is shown in Y, and the superimposition of X and Y is shown in Z. NOR - Novel Object Recognition test; MWM - Morris water maze; SOM+ - Somatostatin-positive interneurons and GAD-67+ - Glutamic acid decarboxylase 67-positive interneurons. Iba-1: Ionized calcium-binding adaptor molecule-1.

### Partial ablation of hilar SOM+ interneurons leads to increased c-fos + staining in DG and CA3

Hilar-perforant-path-associated interneurons (HIPP cells) in the DG hilus are SOM+ cells that distribute axon fibers in the molecular layer, providing dendritic inhibition of DG granule cells[26]. We hypothesized that partial ablation of SOM+ neurons in the DG hilus would thus increase the activity in DG granule cells. Indeed, c-fos staining in DG hilus (p<0.05), DG granule cell layer (GCL) (p<0.05), and total DG (p<0.05) was increased compared to controls, consistent with hyperactivity. c-fos staining in the CA3 subregion of the hippocampus was also increased (p<0.05), whereas c-fos staining in the CA1 subregion of the hippocampus was unchanged (p= 0.9808) (Fig.2).

**Fig. 2.**
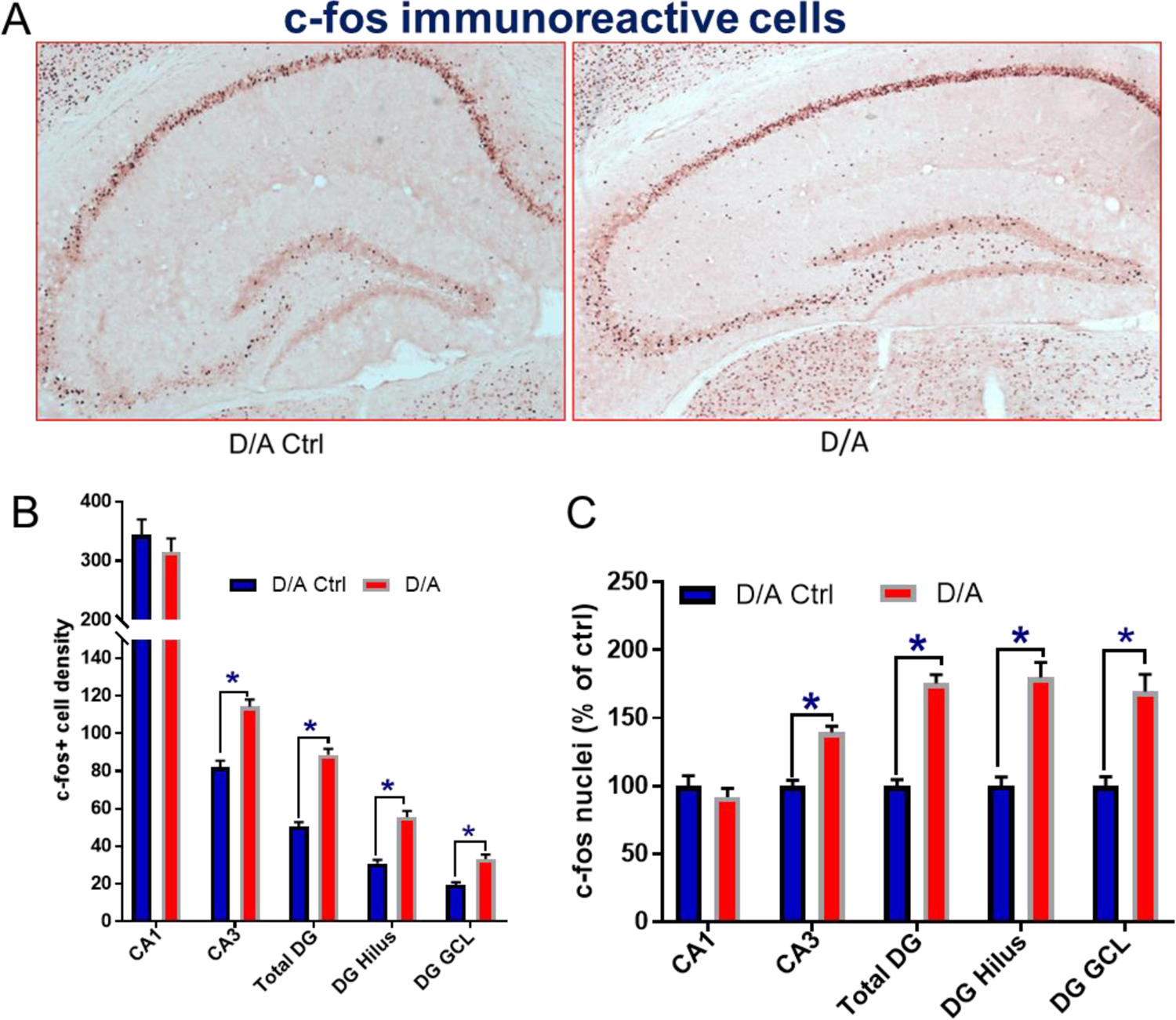
Ablation of Hilar SOM+ interneurons increased c-fos+ neurons in the DG & CA3 regions of the hippocampus. *A)* Representative microscopical images of neurons positive for c-fos. B) Quantification of c-fos-positive cells. C) Histograms show the estimated density of c-fos+ cells in D/A mice expressed as a percentage of D/A control mice. The data were analyzed using NIH Image J software. Statistical analyses (GraphPad Prism) using Two-way ANOVA (interaction between groups and subregions, F (4, 426) = 7.929, p<0.0001; between subregions F (4, 426) = 7.973, p<0.0001; between groups - D/A control and D/A, F (1, 426) = 82.27, p<0.0001, followed by Sidak’s multiple comparisons test showed *p<0.05 vs. D/A Ctrl, (n=4). D/A Ctrl – (Control) injected with an AAV-mCherry virus; D/A-injected with AAV-mCherry-flex-dtA; DG - Dentate gyrus; and GCL-DG granule cell layer.

### Partial ablation of hilar SOM+ interneurons leads to a reduction of the number of SOM+ and GAD-67+ neurons

We then assessed the consequences of the ablation of SOM+ interneurons on the expression of two interneuronal markers. In mice with partial ablation of SOM+ interneurons, the number of SOM+ cells were reduced in DG (p<0.05) and CA3 (p<0.05) compared to controls, but not in CA1 (p= 0.1343) (Fig 3). In addition, the number of GAD-67+ was significantly decreased only in DG hilus of mice with partial ablation of SOM+ interneurons (p<0.01), whereas in CA3 (p= 0.4937) and CA1 (p=0.8552), no significant change was observed for GAD-67+ staining (Fig. 4). These results indicate that our experimental manipulation led to an approximately 50% reduction of SOM+ interneurons in the DG and also in the number of GAD-67+ interneurons.

**Fig. 3.**
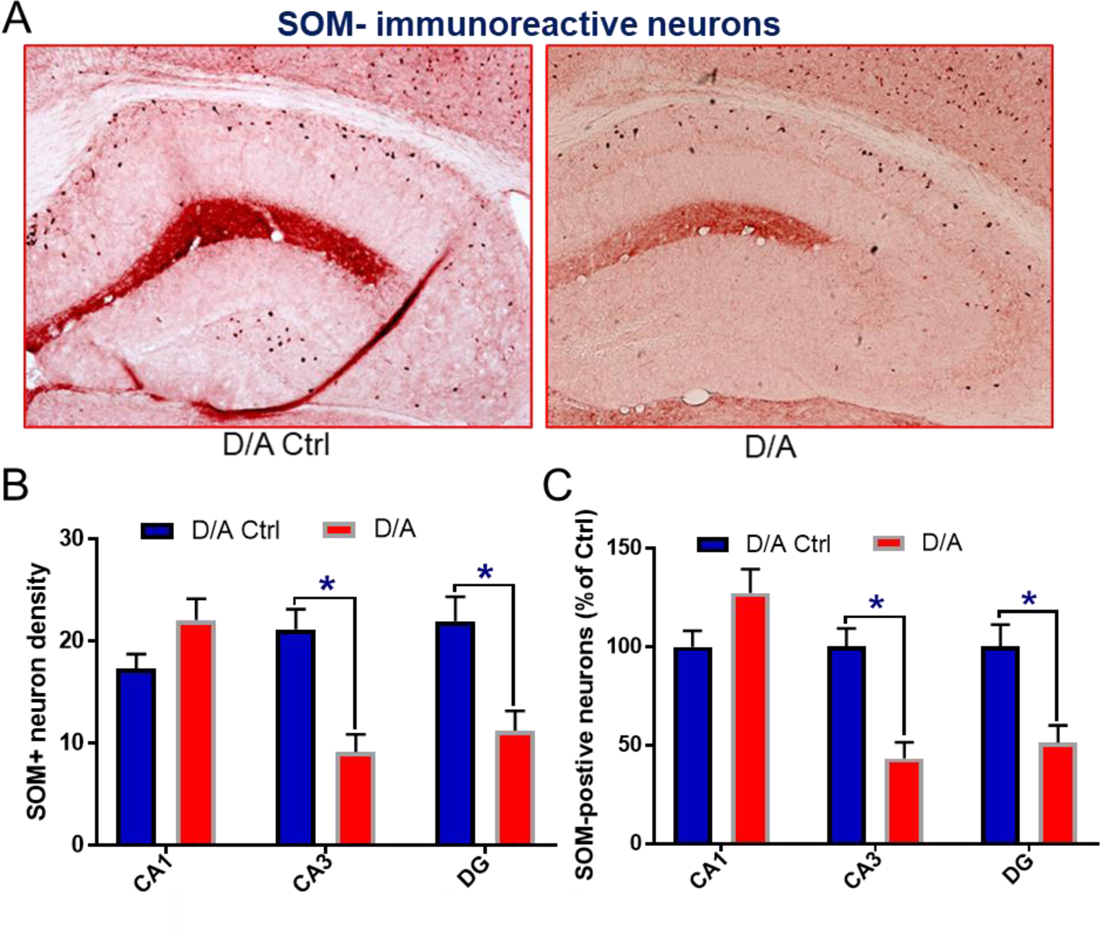
Injection of AAV-mCherry-flex-dtA decreases SOM-immunoreactive neurons in the DG & CA3 region of the hippocampus. *A)* Representative microscopical images of neurons positive for SOM. B) Quantification of SOM-positive neurons. C) Histograms show the estimated density of SOM+ cells in D/A mice expressed as a percentage of D/A control mice. The data were analyzed using NIH Image J software. Statistical analyses (GraphPad Prism) using Two-way ANOVA (interaction between groups and subregions, F (2, 138) = 11.58, p<0.0001; between subregions F (2, 138) = 11.51, p<0.0001; between groups - D/A control and D/A, F (1, 138) = 10.94, p=0.0012, followed by Sidak’s multiple comparisons test showed *p<0.05 vs. D/A Ctrl (n=4).

**Fig. 4.**
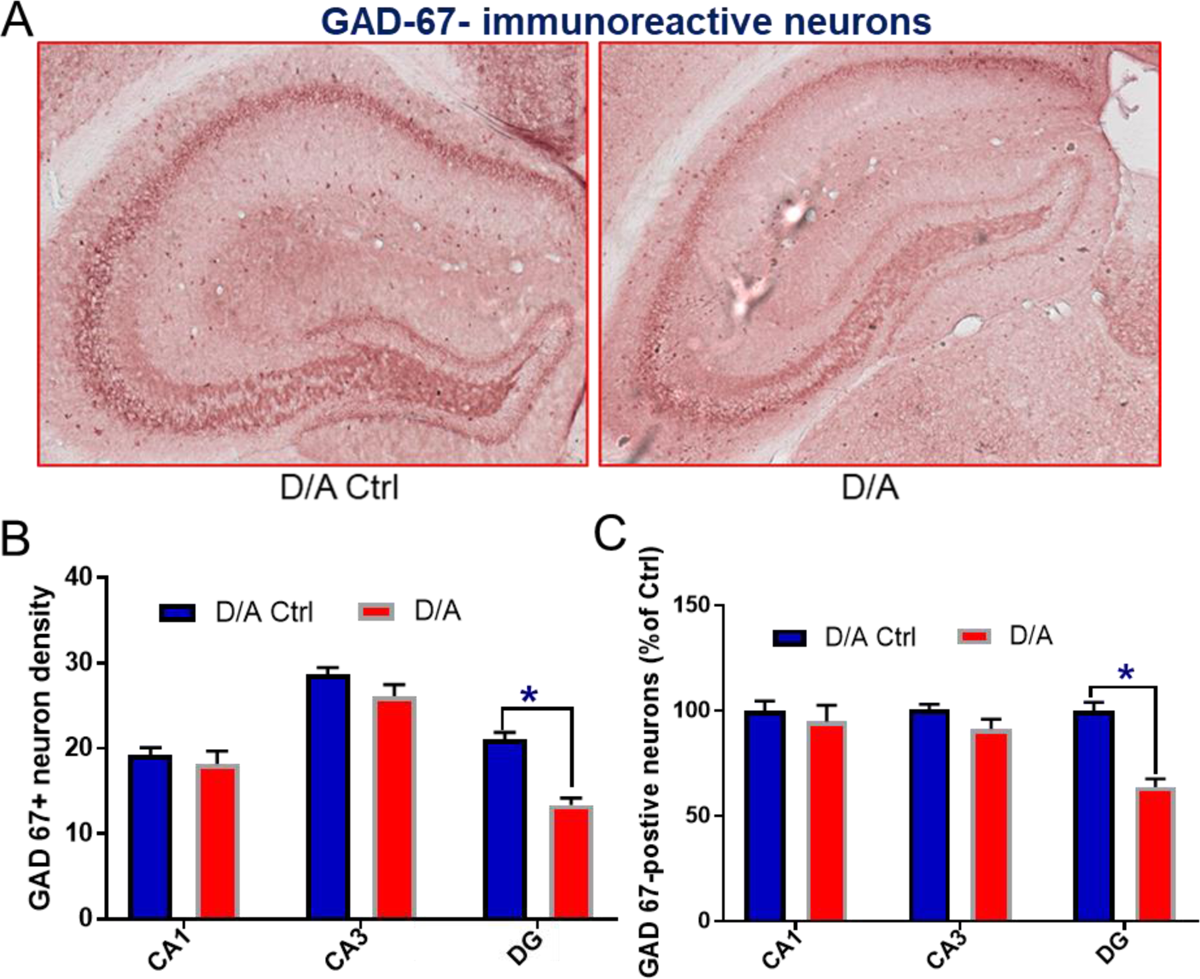
Injection of AAV-mCherry-flex-dtA decreases GAD-67-immunoreactive neurons in the DG region of the hippocampus. *A)* Representative microscopical images of neurons positive for *GAD-67*. B) Quantification of GAD-67-positive neurons. C) Histograms show the estimated density of *GAD-67*+ cells in D/A mice expressed as a percentage of D/A control mice. The data were analyzed using NIH Image J software. Statistical analyses (GraphPad Prism) using Two-way ANOVA (interaction between groups and subregions, F (2, 90) = 5.94, p=0.0038; between subregions F (2, 90) = 6.085, p=0.0033; between groups - D/A control and D/A, F (1, 90) = 17.25, p<0.0001, followed by Sidak’s multiple comparisons test showed *p<0.05 vs. D/A Ctrl (n=4).

### Partial ablation of hilar SOM+ interneurons induces microglial activation in the hippocampus

Microglia activation is a pathological hallmark of neuroinflammation associated with aging and memory impairment, which results in Iba-1 upregulation and an increased cell body size. In the animals injected with the D/A construct, Iba-1 was upregulated in Western blots (hippocampus, p< 0.05, Fig. 5A), and in immunohistochemical studies, the cell body size was increased in the CA3 (p< 0.05, Fig. 5B, C) and DG (p< 0.05, Fig. 5B, C) regions of the hippocampus, but not in the CA1 region of the hippocampus (p =0.1655).

**Fig. 5.**
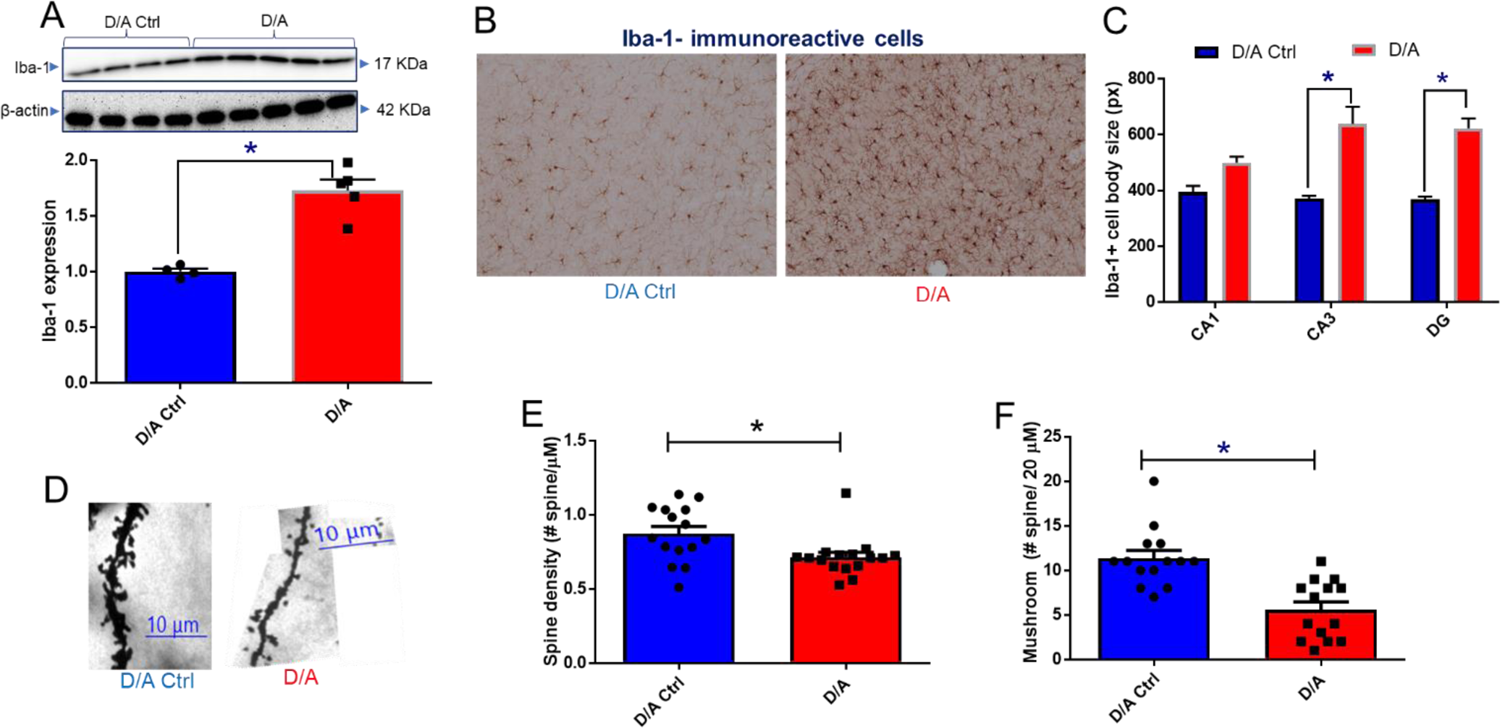
Microglial activation and morphological features of the hippocampus after injection of AAV-mCherry-flex-dtA. A) Representative Western blots images of Iba-1 and β-actin in different groups of mice. *Bar graph images* show mean ± S.E.M. of relative protein expression. The expression was normalized with β-actin. Statistical analyses were performed (GraphPad Prism) using a two-tailed t-test. t_7_=6.342, *\*p<0.05* vs. D/A Ctrl. B) Representable image of Iba-1 expression in microglia cells. C) Histogram showing Iba1-immunostained areas with pixels (px) indicating cell body size. The data were analyzed using N.I.H. Image J software. Statistical analyses (GraphPad Prism) using Two-way ANOVA (interaction between groups and subregions, F (2, 154) = 2.925, p=0.0566; between subregions F (2, 154) = 1.44, p=0.2402; between groups - D/A control and D/A, F (1, 154) = 50.99, p<0.0001, followed by Sidak’s multiple comparisons test showed *p<0.05 vs. D/A Ctrl (n=4). D) Representative pictures of DG dendrite and spine morphologies after D/A injection used for quantification with a 100x objective. E) Spine density in DG. F) Density of mushroom-type spines (width >0.6 μm) in DG. The data were analyzed using RECONSTRUCT software (http://synapses.clm.utexas.edu). Statistical analyses were performed (GraphPad Prism) using a two-tailed t-test (n=4). E) t_29_=2.725, *p<0.05 vs. D/A Ctrl (n=4). F) t_26_=4.643, *p<0.05 vs. D/A Ctrl.

### Partial ablation of hilar SOM+ interneurons leads to decreased hippocampal spine densities

Normal aging is associated with impairments in cognitive function, including memory. These impairments are linked to specific and relatively subtle synaptic alterations in the hippocampus [27].To evaluate the impact of the ablation of hilar SOM+ interneurons on hippocampal spine densities, we analyzed the dendrites of granule cells in the dentate gyrus. D/A injection significantly down-regulated spine density (p<0.05, Fig. 5D, E) and mushroom-type spine density (p< 0.05, Fig.5D, F) in the hippocampus.

### Partial ablation of hilar SOM+ interneurons induces recognition memory impairment in the NOR test

We investigated the effect of partial ablation of SOM+ interneurons on recognition memory in the NOR test. This test is based on the natural tendency of mice to explore a novel object rather than a familiar object. Mice with partial ablation of SOM+ interneurons preferred novel objects over familiar objects to a lesser degree than control mice, which resulted in a significant decrease in the recognition index (p<0.05) compared to controls, indicating that recognition memory is impaired in these mice (Fig. 6A). To determine whether the cellularly defined model (Fig.1A) has any behavioral features of age-related cognitive dysfunction, we have performed behavioral experiments with control mice: young adult (3-4 months) and aged (18-19 months) C57BL/6J wild-type mice. As illustrated in Fig. 6B, old (aged 18-19 months) C57BL/6J wild-type mice displayed a significant difference in recognition index (p<0.05) compared to young adult (3-4 months) controls. We observed no significant differences between males and females in the NOR Test (data not shown).

**Fig. 6.**
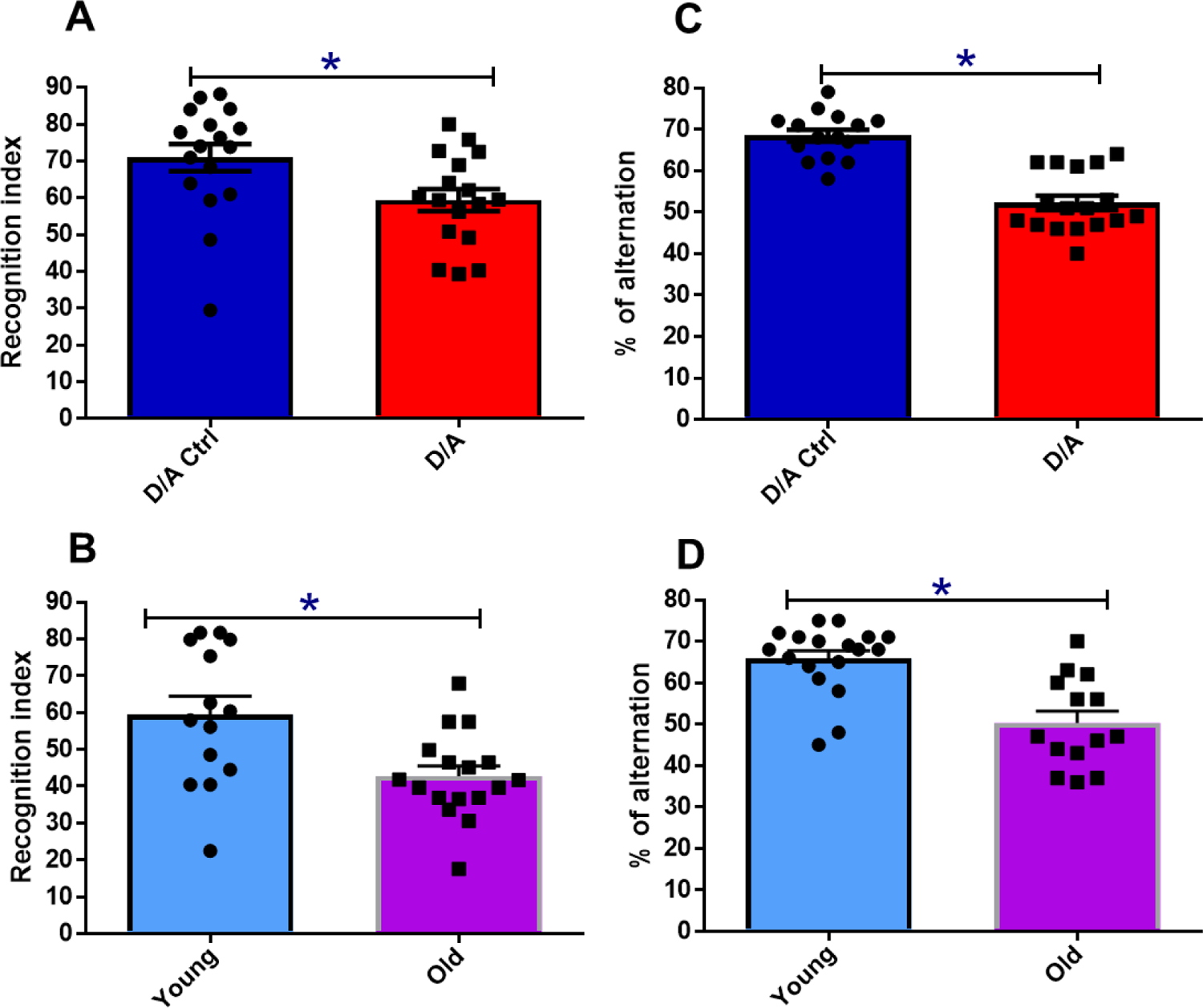
Novel Object Recognition test (NOR) and Y-maze Task in D/A -injected mice and young adults (3-4 months) and aged (18-19 months) C57BL/6J wild-type mice. A &B) Recognition index in the Novel Object Recognition test (NOR) and C&D) Spontaneous arm alternation percentage in the Y-maze test. Results are expressed as mean ± SEM. Statistical analyses were performed (GraphPad Prism) using a two-tailed t-test (n=14-16). A) t_32_=2.426, *p<0.05 vs. D/A Ctrl; B) t_30_=4.594, *p<0.05 young mice. C) t_29_=3.057, *p<0.05 vs. D/A Ctrl; D) t_30_=4.594, *p<0.05 young mice.

### Partial ablation of hilar SOM+ interneurons induces working memory impairment in the Y maze test

The Y-maze task was used to assess short-term working memory. Mice with intact working memory would run in the three arms with spontaneous alternations. We found that mice with partially ablated SOM+ interneurons after D/A injection displayed a reduction in the percentage of correct alternations compared to control mice (p<0.05), indicating that short-term working memory is impaired (Fig.6C). Likewise, aged (18-20 months) C57BL/6J wild type mice displayed a lower percentage of alternations than young adult (3-4 months) C57BL/6J wild-type mice (Fig.6D, p<0.05). We observed no significant differences between males and females in the Y maze (data not shown).

### Partial ablation of hilar SOM+ interneurons induces spatial learning and memory impairment

To investigate potential spatial memory deficits, we performed the Morris Water Maze (MWM). During the learning phase, the D/A control mice displayed a significantly shorter latency to reach the platform on day 4 compared to day 1 (p<0.05), while the partial ablation mice did not show any improvement (p>0.05), which resulted in a difference between the two groups on day 4 (p<0.01) (Fig. 7A). The path length in the control mice was reduced on days 3 and 4 (p<0.05), whereas in partial ablation mice there was no change compared to day 1 (p>0.05), resulting in a difference between the groups on day 4 (p<0.01) (Fig. 7B). In the learning phase probe trials on day 5, the partial ablation group spent a reduced cumulative duration in the platform area compared to controls (p<0.05) (Fig. 7C) and also displayed a reduced frequency to enter the platform position (p<0.05) (Fig. 7D).

**Fig. 7.**
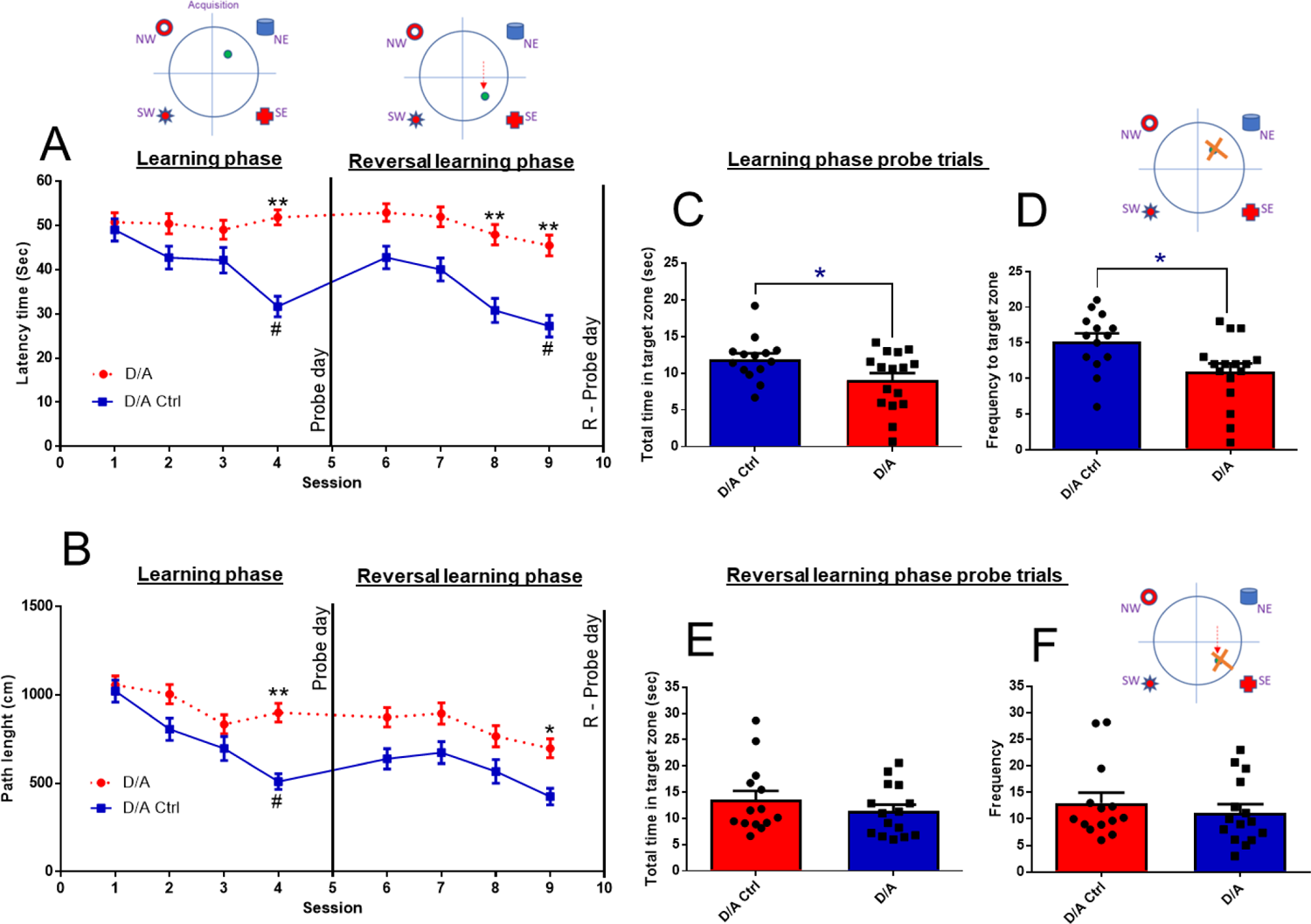
Partial ablation of hilar SOM+ interneurons impaired spatial learning and memory functions in mice. Learning phase Session 1-4 (day1-4); Learning phase probe trials – Session 5 (Day 5); Reversal learning phase Session 6-9 (day 6-9) & Reversal learning probe trials – Session 10 (day 10). A) Latency time (sec) and B) Path length (cm) to the hidden platform in the learning phase (*p<0.05, **p<0.01 vs. D/A Ctrl & #p<0.05 vs. Session 1 - D/A Ctrl) and Reversal learning phase (*p<0.05, **p<0.01 vs. D/A Ctrl & #p<0.05 vs. Session 6 - D/A Ctrl). Statistical analyses were performed (GraphPad Prism) using Two-way ANOVA followed by Sidak’s multiple comparisons test. C) Cumulative duration, and D) Frequency to enter target platform area in learning phase probe trials. E) Cumulative duration and F) frequency to enter the target platform area in the reversal learning phase of probe trials. Statistical analyses were performed (GraphPad Prism) using a two-tailed t-test (n=12-14). C) t_28_=2.223, *p<0.05 vs. D/A Ctrl; D) t_28_=2.630, *p<0.05 vs. D/A Ctrl; E) t_27_=0.982, *p<0.05 vs. D/A Ctrl; F) t_27_=0.736, *p<0.05 vs. D/A Ctrl.

On day 6, the position of the hidden platform was changed to assess reversal learning. In control mice, the latency to find the platform was reduced on day 9 (p<0.05) compared to day 6, whereas there was no change in the latency to find the platform in the partial ablation mice (p>0.05). The latencies of both groups were significantly different on day 8 (p<0.01) and day 9 (p<0.01) (Fig. 7A). In the same trials, the path length of control mice (p>0.05) and partial ablation mice (p>0.05) was unchanged on days 7-9 compared to day 6. However, both groups differed on day 9 with the partial ablation mice displaying an increased path length (p<0.05) compared to the control group (Fig. 7B). In the reversal learning phase probe trials on day 10, both groups of mice did not differ in the cumulative duration spent in the platform area (p>0.05) (Fig. 7E) and in the frequency to enter the platform position (p>0.05) (Fig. 7F).

Further, to assess age-specific differences, we have also performed MWM test experiments with young adult (3-4 months) and aged (18-19 months) C57BL/6J wild-type mice. During the learning phase, the young adult mice displayed a significantly shorter latency to reach the platform on days 3, 5, and 6 compared to day 1 (p<0.05), while the old mice did not show any improvement (p>0.05), which resulted in a difference between the two groups on days 5 and6 (p<0.001) (Fig. 8A). The path length in the young mice was reduced on days 2,3,4,5, and 6 (p<0.05), whereas in old mice there was no change compared to day 1 (p>0.05), resulting in a difference between the groups on day 5 (p<0.01), day 6 (p<0.001) (Fig.7B). In the learning phase probe trials on day 7, the old mice (18-19 months) spent a reduced cumulative duration in the platform area compared to young mice (3-4 months) (p<0.05) (Fig. 8C) and also displayed a reduced frequency to enter the platform position (p<0.05) (Fig. 8D).

**Fig. 8.**
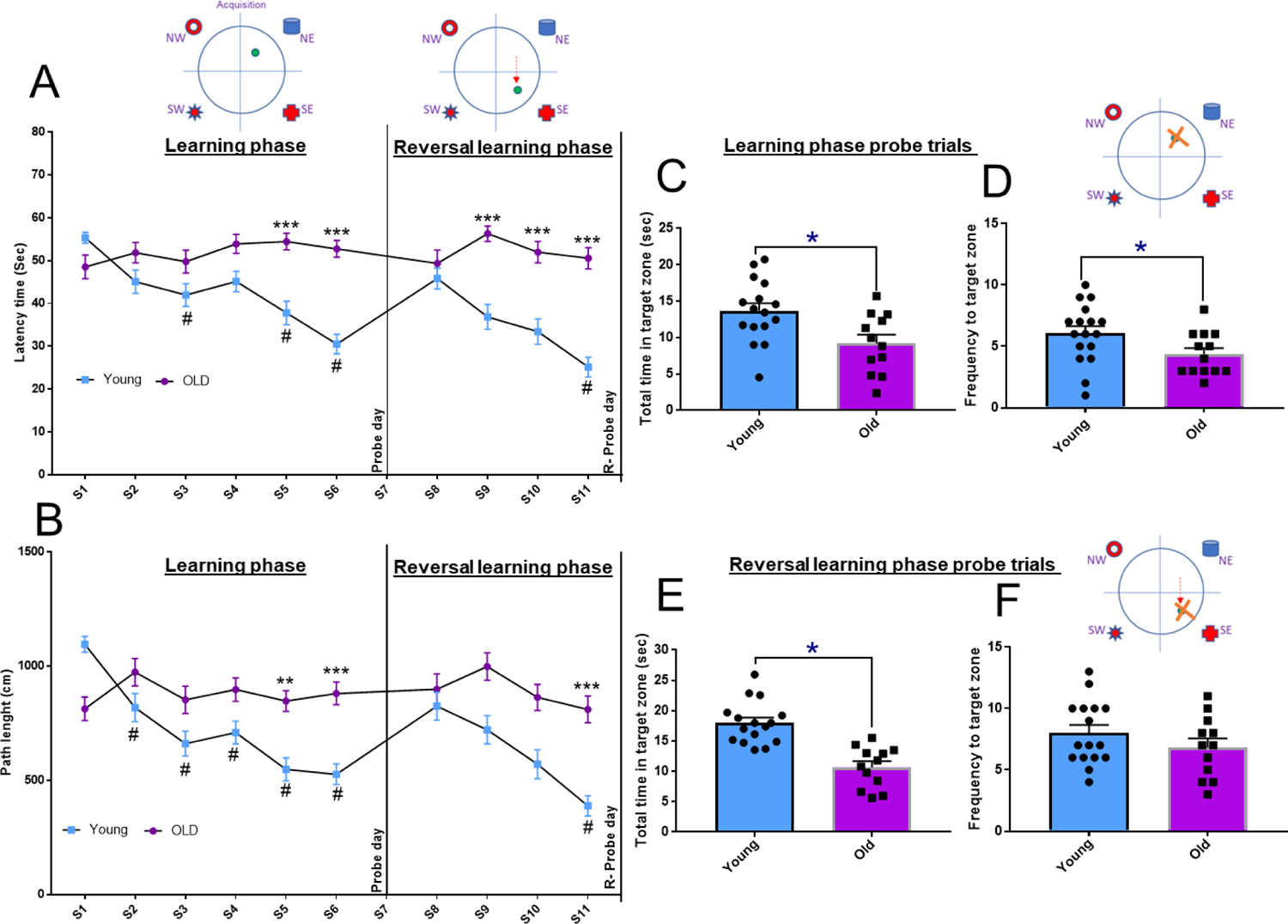
Aging impaired spatial learning and memory functions in C57BL/6J wild-type mice. Learning phase Session 1-6 (day1-6); Learning phase probe trials – Session 7 (Day 7); Reversal learning phase Session 8-11 (day 8-11) & Reversal learning probe trials – Session 12 (day 12). A) Latency time (sec) and B) Path length (cm) to the hidden platform in the learning phase (**p<0.01, ***p<0.001 vs. C57BL/6J wild-type young adult mice (3-4 months) & #p<0.05 vs. Session 1 - Young adult mice) and Reversal learning phase (***p<0.001 vs. young adult mice (3-4 months). Statistical analyses were performed (GraphPad Prism) using Two-way ANOVA followed by Sidak’s multiple comparisons test. C) Cumulative duration and D) Frequency to enter target platform area in learning phase probe trials. E) Cumulative duration and F) frequency to enter the target platform area in the reversal learning phase of probe trials. Statistical analyses were performed (GraphPad Prism) using a two-tailed t-test (n=12-16). C) t_26_=2.772, *p<0.05 vs. *p<0.05 vs. young adult mice (3-4 months); D) t_28_=2.109, *p<0.05 vs. young adult mice (3-4 months); E) t_26_=5.509, *p<0.05 vs. young adult mice (3-4 months); F) t_26_=1.189, *p<0.05 vs. young adult mice (3-4 months).

On day 8, the position of the hidden platform was changed to assess reversal learning. In young mice (3-4 months), the latency to find the platform was reduced on day 11 (p<0.05) compared to day 8, whereas there was no change in the latency to find the platform in the old mice (18-19 months) (p>0.05). The latencies of both groups were significantly different on days 9 (p<0.001), 10 (p<0.001) and 11 (p<0.001) (Fig. 8A). In addition, both groups differed on day 11 with the aged mice displaying an increased path length (p<0.05) compared to the young group (Fig. 8B). In the reversal learning phase probe trials on day 12, both groups of mice did differ in the cumulative duration spent in the platform area (p<0.05) (Fig. 8E) but not in the frequency to enter the platform position (p=0.2452) (Fig. 8F).

Overall, our results indicate that during the learning phase – and to a lesser degree during the reversal learning phase – the partial ablation of SOM+ interneurons in the DG results in cognitive impairments, which is similar to the results in our control experiments with C57BL/6J wild type mice, where aged mice (18-19 months) displayed cognitive impairments in both the learning phase and the reversal learning phase. In other words, the mice with partial ablation of SOM+ interneurons displayed a behavior similar to old mice (18-19 months).

## Discussion

In this study, we investigated the consequences of partial ablation of SOM+ interneurons in the DG hilus for hippocampal learning and memory. We found that our experimental approach led to an approximately 50% reduction of the number of SOM+ interneurons in the DG. This resulted in increased c-fos staining in DG and CA3, and in cognitive impairments in the novel object recognition memory, the Y maze and the MWM with learning and reversal learning phases. Our work shows that partial ablation of DG hilar SOM+ interneurons is sufficient to elicit a phenotype that is remarkably similar to that of aged mice. The mice with partial ablation of DG hilar SOM+ interneurons potentially represent a cellularly defined model of hippocampal aging.

A remarkable finding is that there was an increase in c-fos staining in DG and CA3, as hyperactivity in DG and CA3 are hallmarks of the aging hippocampus. The c-fos gene is a cellular proto-oncogene that can be triggered by neuronal activation, cAMP pathway stimulation, and Ca2+ elevation in the brain [27]. Here, we used c-fos as a marker for neuronal activity in the hippocampus and found increased c-fos expression in the total DG, DG hilus, DG GCL layer and CA3 in the partial ablation mice. Thus, increased c-fos -activity in DG and CA3 is associated with the cognitive impairment of the partial ablation mice. The c-fos staining results in the SOM+ ablation model is similar to the ones that we have previously observed in mice lacking α5-GABA_A_ receptors in dentate granule cells[9], suggesting that a reduction of inhibition of the DG granule cells by hilar SOM+ perforant path-associated (HIPP) neurons[26] may play an important role in this model.

Ablation of SOM+ interneurons in the DG hilus led to impaired performance in the Y maze alternation test, assessing working memory, in the novel object recognition (NOR) test, assessing recognition memory, in initial learning in the Morris water maze (MWM), assessing spatial learning, and in the reversal phase of the MWM, assessing reversal learning. As in our partial ablation mice the experimental manipulation is restricted to the hippocampus, it is reasonable to assume that the behavioral deficits that we see are of hippocampal origin. Based on studies on the role of α5-GABA_A_ receptors in CA1 and DG[8, 9], initial learning in the MWM is largely CA1-dependent and reversal learning largely DG-dependent. Our results suggest that ablation of SOM+ interneurons in the DG results in cognitive deficits that are due to DG and CA1 dysfunction, which are similar to cognitive deficits in aged mice, indicating that a partial ablation of SOM+ interneurons in the dentate hilus is sufficient to explain aging-related cognitive deficits.

The decreased number of SOM-positive and GAD-67-positive interneurons in the hippocampus of partial ablation mice reported here is consistent with findings in other rodent models of aging[28]. Andrews-Zwilling et al. (2010) reported that apoE4 knock-in (KI) mice had an age-dependent decrease in hilar GABAergic interneurons that correlated with the extent of learning and memory deficits, as determined in the Morris water maze in aged mice, which is similar to our findings in mice with partial ablation[29]. Treating apoE4-KI mice with the GABA_A_ receptor potentiator pentobarbital rescued the learning and memory deficits. In neurotoxic apoE4 fragment transgenic mice, hilar GABAergic interneuron loss was even more pronounced and also correlated with the extent of learning and memory deficits[29]. Spiegel et al. (2013) reported that the hilar SOM-positive interneurons exhibit an age-dependent change in numbers in outbred Long-Evans rats, with significantly reduced numbers in the subpopulation of aged rats with behavioral impairment while unimpaired aged rats had SOM+ interneuron numbers that did not differ from young adults[15]. Interestingly, postmortem studies of patients with AD have shown reduced SOM expression in the hippocampus and cortical areas[30–32], and the decreased expression was correlated with the severity of cognitive deficits[33] and neuropathology[34]. The current findings of memory impairments in our partial ablation mice are therefore consistent with other evidence in animal models and studies in humans indicating that a loss of hilar integrity may be underlying such an age-related memory impairment. Our findings are also consistent with a potential contribution of GABAergic interneuron impairment to the pathogenesis of AD.

It is important to note that all SOM-expressing hilar interneurons colocalize with GAD-67 [35, 36]. The current data show that the decline in SOM-expressing hilar interneurons accounted almost entirely for the magnitude of hilar GAD67-positive neuron loss observed in partial ablation mice. This strongly suggests that hilar vulnerability promoting cognitive impairment during aging predominantly involves the GABAergic hilar interneurons that are also positive for SOM. Decreased GAD-67 and SOM protein expression may impact hilar interneuron function in partial ablation mice. While other approaches would be needed to fully document the physiological consequences of hilar interneurons lacking SOM and GAD-67 expression, evidence suggests that reduced levels of GAD-67 protein expression would result in diminished functional inhibition. In support of this possibility, Lau and Murthy (2012) reported that decreased levels of cytosolic GAD-67 expression are associated with reduced GABA synthesis, diminished GABA release at the synapse, and decreased miniature inhibitory postsynaptic current amplitudes (mIPSCs) in cultured hippocampal neurons[37]. The concept that reduced inhibitory function in the hippocampal memory network contributes to cognitive dysfunction, as suggested by our findings, is consistent with the presence of hyperactivity of the DG and CA3 subregions in both humans and animal models with age-related memory impairment[16, 38, 39]. Moreover, targeting that hyperactivity has been found to improve hippocampal-dependent memory function[29, 38, 40].

In the current study, we found that decreased SOM-positive and GAD-67-positive interneurons in the hippocampus of partial ablation mice leads to microglial activation. Moreover, we found that neuroinflammation impaired hippocampal spine density, leading to cognitive impairment, corroborating previous studies[28, 41]. Further, Mahmmoud et al (2015) report that the decrease in spine density and mushroom spines (which have been proposed to be “memory spines”) in the dentate gyrus is linked to spatial and working memory [42]. In the present study, D/A injected memory-impaired partial ablation mice showed significantly reduced dendritic spine density and mushroom spines in the dentate gyrus, in line with the role of mushroom spines as “memory spines.”

We have recently reported on a model of hippocampal aging based on chronic chemogenetic inhibition of SOM+ interneurons in the DG [28]. The partial ablation model displays a few notable differences to this chemogenetic model. Whereas in the chemogenetic model increased c-fos staining is only observed in DG, in the ablation model this is present in both DG and CA3. Likewise, while in the chemogenetic model the SOM+ neuron density is reduced only in DG, in the ablation model it is reduced in both DG and CA3. On the other hand, microglial activation has been observed in DG, CA3 and CA1 in the chemogenetic model, but only in DG and CA3 in the ablation model. However, the degree of microglial activation appears to be greater in the ablation model. The ablation model led to the mice spending less time in the target zone in the learning probe trial in the water maze, which was not seen in the chemogenetic model, whereas in the chemogenetic model mice spent less time in the target zone in the reversal learning probe trial, which was not observed in the ablation model. Overall, it appears that the ablation model may have a more pronounced phenotype than the chemogenetic model, in particular that hyperactivity extends beyond the DG to the CA3, as has been observed in aging rodents. Thus, the ablation model has a few features that may make it preferable to the chemogenetic model for several types of future studies.

In conclusion, our results show that the partial ablation of hilar SOM-positive interneurons is sufficient to induce changes similar to those observed in aged rodents, such as reduction of the SOM and GAD-67 markers in the DG hilus, hyperactivity in DG and CA3 of the hippocampus and cognitive impairments. This suggests that the changes elicited by partial ablation of hilar SOM-positive interneurons may contribute to the development of cognitive dysfunction in aging animals. Furthermore, mice with a partial ablation of hilar SOM+ interneurons may represent a model for studying mechanisms underlying cognitive decline in aging.

## Acknowledgements

We thank Dr. CheMyong Ko for providing access to his microscopes.

## Author contributions

Conceptualization: UR; Investigation: RN; Analysis and Interpretation: RN, JL,UR; Methodology: RN,JL, MK, MW, CCH, CDC, UR; Funding acquisition: UR; Supervision: UR; Writing – original draft: RN, UR; Writing – review & editing: RN,JL, MK, MW, CDC,CCH,UR.

## Data availability

All original research data will be archived in the Harvard Dataverse repository.

## Ethics approval

Permission for animal experiments were obtained from the University of Illinois IACUC.

## Consent to participate

N/A.

## Consent to publish

N/A.

